# Collective cell migration and residual stress accumulation: modeling consideration

**DOI:** 10.1101/2019.12.18.881144

**Authors:** I. Pajic-Lijakovic, M. Milivojevic

## Abstract

Stress generation during collective cell migration represents one of the key factors which influence the configuration of migrating cells, viscoelasticity of multicellular systems and their inter-relation. Local generation of stress (normal and shear) is significant even in 2D (up to ~100 − 150 *Pa*). Compressive stress is primarily accumulated (1) within a core region of migrating cell clusters during their movement through the dense environment and (2) during the collisions of migrating cell clusters caused by uncorrelated motility. Shear stress can be significant within perturbed boundary layers around migrating clusters. Cells are more sensitive to the action of shear stress compared with compressive stress. Shear stress of a few *Pa* significantly influences cell state. Deeper insight into cell strategy to minimize undesirable shear stress is a priority in order to understand various biological processes such as morphogenesis, wound healing and cancer invasion. We pointed out to cause-consequence relations of these complex phenomena based on rheological modeling consideration in order to stimulate further experimental work.

Cell strategy should be connected with the type and distribution of adhesion contacts such as adherens junctions and tight junctions per migrating clusters in order to (1) reinforce the cluster structure perpendicular to the direction of cell migration and (2) ensure structural elasticity of cluster in the direction of migration. These conditions lead to the stiffness inhomogeneity per single migrating clusters. Cell strategy should also be related to the state of the perturbed boundary layer around the cluster in the context of its thickness and slip effects.

**Statement of Significance:** Collective cell migration induces a local generation of stress (normal and shear) significant even in 2D (up to *100-150 Pa*). Cells well tolerate compressive stress up to a few *kPa*. However, shear stress of a few *Pa* can induce severe damage to vimentin and keratin intermediate filament networks during 1 h, while shear stress of ~60 *Pa* can cause the inflammation in epithelial cells during 5.5 h. Deeper insight into cell strategy to minimize undesirable shear stress is a priority in order to understand various biological processes. Cell strategy should be connected with the type and distribution of adhesion contacts such as adherens junctions and tight junctions per migrating clusters and surrounding perturbed boundary layers.

## 1. Introduction

A more comprehensive account of main features of: (1) collective cell migration, (2) its influence to viscoelasticity of multicellular systems, and (3) feedback impact of viscoelasticity to cell migration are an essential for a wide range of biological processes such as embryo morphogenesis, wound healing, regeneration, and also in pathological conditions, such as cancer (1–6). Recent studies suggest that the dynamic tuning of the viscoelasticity within a migratory cluster of cells, and the adequate elastic properties of its surrounding tissues, are essential to allow efficient collective cell migration (2–3). Consequently, the viscoelasticity depends on (1) viscoelasticity of migrating cell clusters, (2) viscoelasticity of surrounding resting cells, and (3) configuration of migrating cells (4,7).

One of the key control parameters which influence the configuration of migrating cells and the rate of its change is the residual stress accumulation as a consequence of collective cell migration (4). The stress can be normal (compressive and tensile) and shear. Compressive stress is primarily accumulated within a core region of migrating cell clusters during their movement through the dense environment (4) and during collisions of migrating cell clusters caused by uncorrelated motility (8,9). Collision of velocity fronts can lead to jamming state transition (10,11). This stress accumulation can suppress cell migration and consequently influences the lifetime of migrating clusters and its distribution depending on: (1) magnitude, (2) duration, and (3) a way of locally generated stress. Shear stress could be significant within perturbed boundary layers around migrating clusters.

Residual stress accumulation is caused by intrinsic and extrinsic cellular processes. Intrinsic processes are: (1) gene expression differences, (2) cell signalling and (3) their inter-relations. The coordinated movement of clusters of cells with respect to the surrounding tissue is often guided by short- or long-range signaling (1). Cells communicate with each other via direct contact (juxtacrine signalling), over short distances (paracrine signalling), or over large distances (endocrine signalling). Gene expression induces time delay in cell response to various mechanical and biochemical stimulus (6). This time delay might be relevant for cell coupling because what cells acquire at the present time is the information of surrounding cells some time ago. These perturbations can induce that (1) cells in the same population respond to different signals and/or (2) cells behave differently in response to the same signals (1) that could lead to uncorrelated motility. The uncorrelated motility caused by a collision of velocity fronts is significant even in 2D (9). Consequently, the collision of velocity fronts can induce the formation of stagnant zones that leads to local generation of compressive and shear stresses within multicellular systems and the change of the configuration of migrating cells. Extrinsic processes depend on substrate mechanical properties and/or neighboring tissue forces (1). These processes are additionally influenced by significant stiffness difference between migrating cell clusters and surrounding resting cells (12). Cumulative effects of these mechanical and biochemical processes influence the configuration of migrating cells and the rate of its changes and on that base the viscoelasticity of multicellular systems on various space and time scales (4,5).

Tambe et al. (8) considered stress generation within collective migrated epithelial cell monolayers. They reported that local traction must be balanced by local gradients in monolayer normal and shear stresses. They estimated stresses values up to 100-150 *Pa*. Du Roure et al. (13) reported that traction stress per single cells is 2.5-3.8 *kPa* while the maximal traction stress inside the epithelium is ~12.7 *kPa*. This result is expectable in the context of the fact that cells are able to generate the stress of 1 *kPa* per single adherens junction (AJs) between two contractile cells (14). However, the residual stress generation could be more intensive in 3D systems (4).

For a deeper understanding of collective cell migration and its influence to viscoelasticity, it is necessary to estimate separately cells response under shear and compressive stresses. Flitney et al. (15) pointed out that some cells, most notably by endothelial cells, well tolerate shear stress up to a few *Pa*. Pitenis et al. (16) pointed out that higher shear stress ~60 *Pa* is sufficient to induce inflammation in epithelial cells during time period of 5.5 h. Interestingly, cells also well tolerate much higher values of compressive stress compared with shear stress, i.e. up to a few *kPa* (17–19). Motivated by these recent findings we pointed out the main causes of local stress accumulation caused by collective cell migration and discussed (1) cell response under shear and compressive stresses, as well as (2) cells strategy to minimize undesirable generation of shear stress based on the rheological modeling consideration. Cell strategy is connected with distribution of adhesion contacts such as adheren junctions (AJs) and tight junctions (TJs) per single migrating cluster and within surrounding perturbed boundary layer. AJs are cadherin-catenin complexes linked to actin filaments. TJs are high affinity complexes formed by transmembrane proteins, including claudins, occludins and tricellulins associate with numerous peripheral proteins. These complexes are also linked to actin filaments. AJs form relatively weak adhesion contacts compared with TJs. Both types of cell-cell adhesion contacts significantly influence cell polarization, signalling and migration (20). This consideration can be useful for biologists to plane their experiments for considering stress distribution during 3D collective cell migration and its influence on configuration of migrating cells and the distribution of adhesion contacts.

## 2. Stress accumulation within a migrating cell cluster

Deeper insight into causes of the stress accumulation during collective cell migration from the rheological standpoint is prerequisite for understanding underlying mechanisms of cells adaptation under various stress conditions and cell strategy to minimize generated stress. Consequently, we considered the stress accumulation: (1) within migrating cell cluster during its movement through a dense environment made by resting cells and (2) during a collision of migrating cell clusters caused by uncorrelated motility. Following aspects of collective cell migration motivated this modeling consideration:

1. Active (contractile) cells are much stiffer than the passive ones due to the accumulation of contractile energy. Lange and Fabry (12) reported that muscle cells can change their elastic modulus by over one order of magnitude from less than 10 *kPa* in a relaxed (resting) state to around 200 *kPa* in a fully activated (contractile) state. This interesting property of cellular dynamics motivated us to develop the so-called “pseudo-blend model” by the analogy with physics for describing long-time rearrangement of cells (4,7).
2. Collectively migrated cells generate viscoelastic waves (9). These waves can be related to the relaxation ability of multicellular systems in the context of successive relaxation cycles which leads to distribution and accumulation of the residual stress (4,21).
3. During migration, cell clusters behave as a viscoelastic solid (22). All cells within the migrating group move, maintaining cell–cell adhesions (23,24). Shellard and Mayor (25) reported that larger migrated clusters behave as supra-cells by establishing supracellular cytoskeletal organization. They experimentally determined two regions: (1) ordered region of cluster and (2) surrounding perturbed boundary layers. Circular movement of cells is observed within the perturbed layers.
4. Mechanosensitivity of E-cadherin turnover depends on p120-catenin, a protein that binds to the E-cadherin tail and blocks access to the endocytic machinery. P120 is released from AJs when stress is high and re-associates with junctions when stress relaxes. Under high-stress, p120-catenin complexes are released into the cytoplasm, destabilizing E-cadherin complexes and lead to an increase the E-cadherin turnover (26).
5. Malinova and Huvanees (27) considered various states of AJs depending on cell active/passive states. AJ between two passive cells corresponds to linear adhesion junction. However, the AJ between two active/contractile cells corresponds to symmetric focal adhesion junction while the AJ between two migrating cells changes to asymmetric focal adhesion junction. This asymmetry can be related to the stress increase and its distribution at the cell-cell AJs caused by cell migration.

### 2.1 Stress accumulation within migrating cell cluster during its movement through dense environment

The main cause of local stress accumulation per single migrating cell cluster during its sliding through dense surrounding is the dynamics at the biointerface between migrating cell cluster and surrounding perturbed boundary layer made by migrating and resting cells (4). The biointerface is expressed as: ℜ_*i*_ = ℜ_*i*_(*x*, *y*, *z*). The change of the biointerface state occurs during multi time relaxation cycles. Step displacements of the migrating cell cluster induce the generation of local compressive and shear strains within migrating and resting cell “pseudo-phases” at the interface. These displacements depend on translation, rotation and random speeds of the cluster (4) and the number density and type of adhesion contacts (20). Local strains lead to a generation of local stresses and their relaxations. These stresses relaxations correspond to short-time relaxation cycles ∆*t* equal to 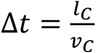 (where *v*_*C*_ is the average speed of migrating cell cluster and *l*_*C*_ is the average size of a single cell). Clark and Vignjajevic (28) reported that the average speed of migrating cell clusters during embryogenesis is about 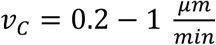. For the average cell size *l*_*C*_ ≈ 10 *μm* the duration of short-time relaxation cycle is equal to ∆*t* ≈ 10 − 50 *min*. Consequently, the duration of single relaxation cycle is ∆*t* ≤ *t*_*p*_, where *t*_*p*_ is the cell persistence time. The order of magnitude of the cell persistence time is a several tens of minutes (29). Residual stress accumulation influences (1) the state of single cells and configuration of migrating cells in the context of migrating-to-resting cell state transitions (4,11), (2) the state of cell-cell adhesion contacts (26) and (3) their inter-relations (27). Cell cytoskeleton is expected to remodel by actin polymerization and de-polymerization while migrating, which can contribute to stress relaxation (30). Many short-time relaxation cycles occur during long-time relaxation cycle. Long-time relaxation cycle corresponds to the contact time *τ* between migrating cell cluster and surrounding tissue equal to 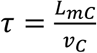 (where *L*_*mC*_ is the effective diameter of migrating cell cluster which is an order of magnitude higher than the size of single cells (3,25). Consequently, the contact time corresponds to a time period of a few hours. The contact time is *τ* ≤ *t*_*lt*_ (where *t*_*lt*_ is the lifetime of migrating cell cluster). The lifetime of migrating clusters should be correlated with the residual stress accumulation.

We will consider three subsystems (1) migrating cell cluster, (2) perturbed boundary layer around the cluster, and (3) surrounding unperturbed resting cells. For 3D modeling consideration, we introduced the space coordinate ℜ = ℜ(*x*, *y*, *z*). All subsystems are characterized by the standpoint of rheology. Cause-consequence relations between: (1) type and number density of adhesion contacts, (2) displacement field, (3) induced strains, and (4) generated stresses which lead to accumulation of strain energy density within a migrating cell cluster and the surrounding boundary layer through many short-time relaxation cycles is presented schematically in Figure 1. These inter-relations and corresponding negative feedback controls describe the interaction of two viscoelastic subsystems such as migrating cell cluster and perturbed boundary layer of surrounding tissue.

**Figure 1.**
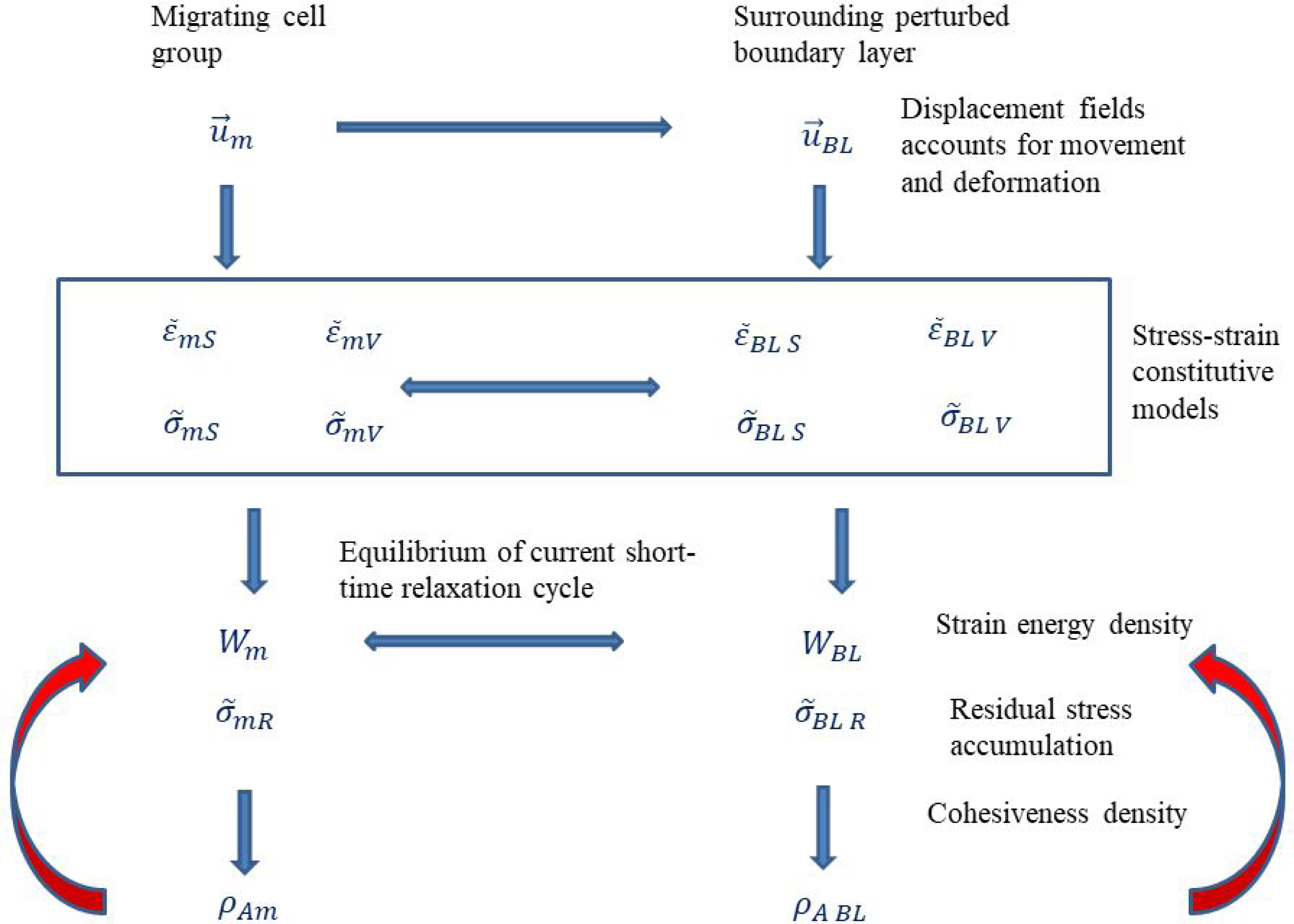
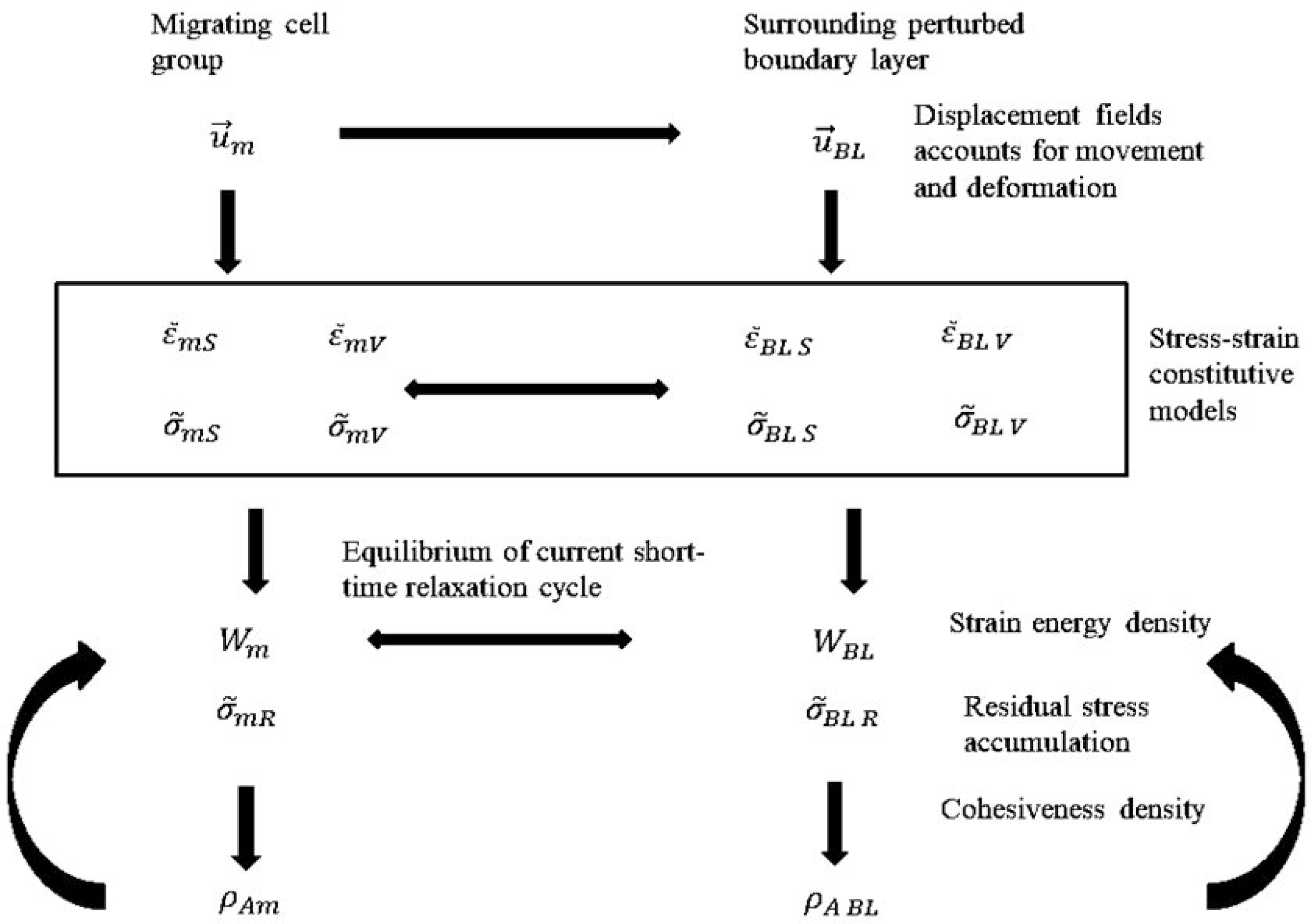
Schematic presentation of the cause –consequence relation between residual stress generation and changes of the cohesiveness density within a migrating cluster and perturbed boundary layer.

All cells within the migrating cluster move, maintaining cell–cell adhesions (24). The stiffness of the migrating cell cluster is influenced by (1) stiffness of migrating cells themselves and (2) type and number density of cell-cell adhesion contacts.

After single short-time relaxation cycle the corresponding current equilibrium state (*t*_*eq*_, *τ*) at the biointerface can be expressed in the context of the strain energy density as:

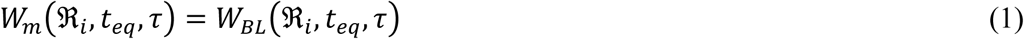

where *W*_*m*_(ℜ_*i*_, *t*_*eq*_, *τ*) is the strain energy of migrating cluster while *W*_*BL*_(ℜ_*i*_, *t*_*eq*_, *τ*) is the strain energy of the boundary layer at the biointerface. The strain energy *W*_*m*_(ℜ_*i*_, *t*_*eq*_, *τ*) is expressed as:

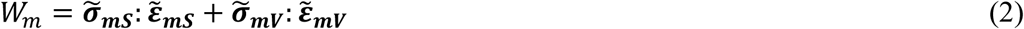

where 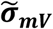 is the volumetric (compressive and/or tensile) stress of migrating cells, 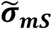 is the shear stress of migrating cells, and 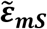 is the shear strain of migrating cells, and 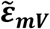 is the volumetric strain of migrating cells. Volumetric and shear stresses consist of elastic and viscous contributions and can be expressed as: 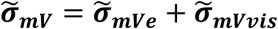 for volumetric stress and 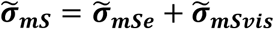 for shear stress (where 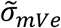 and 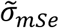 are elastic contributions while 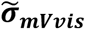 and 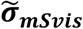 are viscous contributions, respectively), similarly as was formulated by Murray et al. for viscoelastic systems (31). The strain energy *W*_*BL*_(ℜ_*i*_, *t*_*eq*_, *τ*) is expressed as:

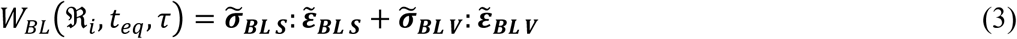

where 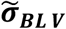 is the volumetric (compressive and tensile) stress of the boundary layer at the biointerface, 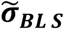 is the shear stress, and 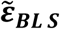 is the shear strain, and 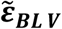 is the volumetric strain of the boundary layer at the biointerface. Volumetric and shear stresses consist of elastic and viscous contributions and can be expressed as: 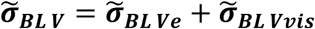 for volumetric stress and 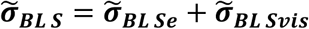 for shear stress (where 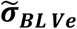 and 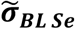 are elastic contributions while 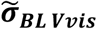 and 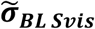 are viscous contributions, respectively).

The corresponding volumetric and shear strains of the migrating cluster at the biointerface are equal to

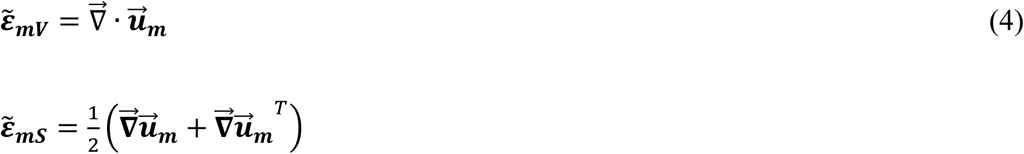

where 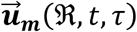 is the local displacement field of migrating cell cluster at the biointerface (4):

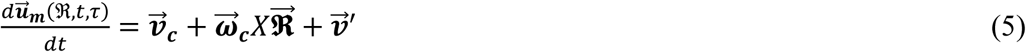

where 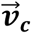 is the translation speed, 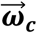 is the angular speed and 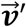 is the random speed.

The corresponding volumetric and shear strains of the boundary layer at the biointerface are equal to:

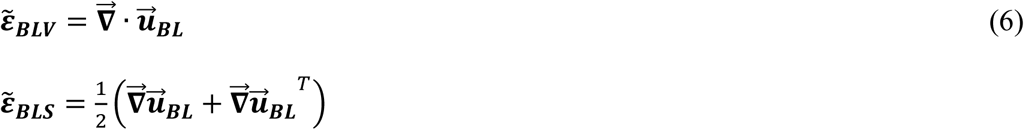

where 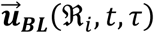 is the local displacement of the boundary layer at the biointerface. Displacement rate of migrating cell cluster at the biointerface 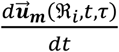 induces displacement of perturbed boundary layer (i.e. the subsystem 2) 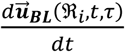. Pajic-Lijakovic and Milivojevic (4) related displacement changes of 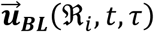 and 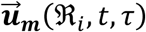 at the biointerface based on the thermodynamical approach at mesoscopic level. It was expressed as:

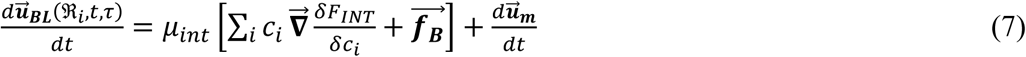

where *μ*_*int*_ is the mobility of the biointerface, 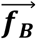 is the Brownian random force, *c*_*i*_ is the concentration of the i-th signalling molecule *F*_*INT*_ is the free energy functional at the biointerface which accounts for cell signalling in order to adapt of surrounding tissue on cell movement.

The momentum balance at the interface at the end of current short-time relaxation cycle for the time set (*t*_*eq*_, *τ*) has been expressed as (32):

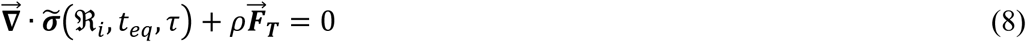

where 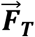 is the average traction force of migrating cells near the biointerface, *ρ* is the average density of cell cluster, and 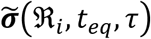 is the total residual stress equal to:

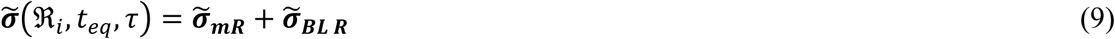

where 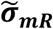 is the residual stress of migrating cells at the interface, 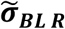 is the residual stress of the boundary layer at the interface. Momentum balance expressed by eq. 8 indicates that stress of migrating cells at the biointerface is a product of cumulative effects of cell tractions and the generated stress within the surrounding boundary layer.

The current equilibrium state should satisfy additional condition for whole migrating cell cluster and surrounding boundary layer for ℜ = ℜ(*x*, *y*, *z*) expressed as

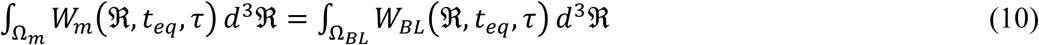

where Ω_*m*_ is the volume of migrating cluster and Ω_*BL*_ is the volume of surrounding boundary layer. The maximum strain energy density is generated within the core region of the migrating cell cluster and decreases up to ℜ = ℜ_*i*_. If the average strain energy density within the core region reaches the threshold value *W*_*c*_, cell migration can be suppressed. The time period for 〈*W*_*m*_〉_*core*_ → *W*_*c*_ corresponds to the lifetime of the migrating cell cluster. The value of *W*_*c*_ hasn’t been experimentally determined. On the other hand, the maximum strain energy density within a perturbed boundary layer is obtained at the interface and decreases up to the unperturbed cell parts. The volume fraction of moving cells within a migrating cell cluster is ϕ_*m*_(ℜ, *t*, *τ*) → 1. Perturbations of cell state within a boundary layer around the migrating cell cluster are induced: (1) biochemically by signalling of migrating cells from one side of the boundary layer and resting cells from the other side and (2) mechanically by a generation of the compressive and shear stresses caused by moving of migrating cell cluster (5). Consequently, single cells within the boundary layer at the same time can receive the signalling molecules from migrating cells and from resting cells. These perturbations can induce that (1) cells in the same population respond to different signals and/or (2) cells behave differently in response to the same signals (1). These phenomena could lead to disordered single cell migration from time to time within the boundary layer in order to (1) decrease slip effects and (2) ensure tissue continuity. Consequently, the volume fraction of moving cells within a boundary layer decreases from ϕ_*m*_ = 1 to ϕ_*m*_ = 0 at the boundary between perturbed layer and surrounding resting cells.

#### 2.1.1 Residual stress accumulation during successive relaxation cycles

The prerequisite of cells to preserve their biological function is the ability of stress relaxation. During relaxation stress decreases from initial value to the equilibrium value for both, migrating cell cluster and surrounding perturbed boundary layer. The stress relaxation time *τ*_*R*_ should be lower than the duration of a single short-time relaxation cycle, i.e. *τ*_*R*_ < ∆*t*.

Distribution of normal (compressive and tensile) and shear stresses has been measured for 2D systems such as monolayer sheets under *in vitro* conditions (8). They determined the instantaneous stress distribution rather than its time-change. Consequently, stress relaxation phenomena haven’t been reconstructed from their data. Stress distribution within 3D multicellular systems caused by collective cell migration under *in vivo* conditions hasn’t been measured yet. Marmottant et al. (33) considered stress relaxation of cellular aggregate under constant strain (i.e. the aggregate shape) condition caused by the aggregate uni-axial compression between parallel plates. The long-time rheological response of cellular aggregate occurs via collective cell migration (34,35). The stress decreases exponentially with the relaxation time equal to 3-14 *min* up to equilibrium value (33). Stress relaxes from ~27 Pa to the residual stress value equal to ~17 Pa during 25 min. This time period corresponds to the duration of single short-time relaxation cycle. The phenomenon of stress relaxation has been related to the adaptation of adhesion contacts (23).

We described the stress relaxation under constant strain condition and corresponding residual stress accumulation from the rheological standpoint for a simple linear model such as the Zener model proposed for viscoelastic solid. This model introduces one relaxation time for stress under constant strain condition *τ*_*R*_ and the other relaxation time for strain under constant stress condition 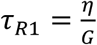 (where *η* is the viscosity and *G* is the elastic modulus). This model corresponds to the rheological response of cell aggregate under uniaxial compression between parallel plates in the context of (1) stress relaxation under constant aggregate shape conditions, (2) aggregate shape relaxation under constant stress conditions, and (3) aggregate free relaxation (its rounding) measured by Mombash et al (36) and Marmottant et al. (33). The Zener model can be expressed for the k-th short-time cycle as:

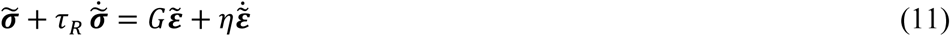

where 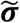 is the local shear or normal stress equal to 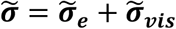, 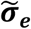 is the reversible –elastic contribution to stress, 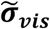 is the irreversible –viscous contribution to stress, 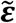 is the corresponding strain, *G* is the local elastic modulus, *η* is the local viscosity, *τ*_*R*_ is the stress relaxation time, 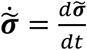 and 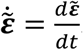. Marmottant et al. (33) determined the effective viscosity of multicellular epithelial aggregate equal to *η* = 4.4*x*10^5^*Pas*. The relaxation of local stress under constant strain per cycle 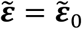 for the k-th cycle can be expressed as:

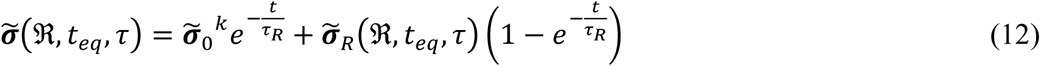

where 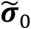 is the initial value of the stress and the residual stress 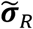 is equal to:

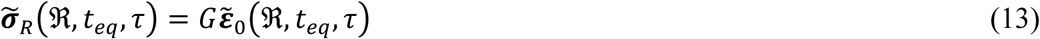

where the corresponding local elastic modulus *G* = *G*(ℜ, *τ*) is constant per single short-time cycle but changes from cycle to cycle. The residual stress increases from cycle to cycle during cluster movement. Local changes of the elastic modulus and viscosity within (1) migrating cluster and (2) surrounding boundary layer quantified the stiffness in-homogeneity and should be correlated with the distribution of cell-cell adhesion contacts. Consequently, the elastic modulus can be expressed for the k-th cycle as:

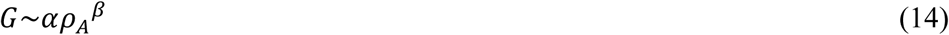

where *ρ*_*A*_ is the number density of adhesion contacts, *α* is the specific energy density per single migrating/resting state, and *β* is the scaling exponent. Both parameters depend on the type of strain, i.e. normal or shear. A similar relation between elastic modulus and cohesive contact density has been formulated by Gaume et al. (37), for granular material. Cellular systems are much complex, but some general principles in the form of this scaling low could be applied at this level of consideration. Local viscosity *η*(ℜ, *τ*) can be expressed as *η* = *Gτ*_*R*1_ (where *τ*_*R*1_ is equal to 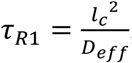, *l*_*c*_ is the average size of single cell, and *D*_*eff*_ is the effective diffusion coefficient). The effective diffusion coefficient is equal to *D*_*eff*_~*k*_*B*_*T*_*eff*_*μ*_*int*_ (where *k*_*B*_ is Boltzmann constant, *T*_*eff*_ is the effective temperature and *μ*_*int*_ is the mobility of the biointerface). The concept of effective temperature has been used for considering rearrangement of various thermodynamical systems (near to equilibrium and far from equilibrium) from glasses and sheared fluids to granular systems (38). Pajic-Lijakovic and Milivojevic (5,11) used this concept to describe cell rearrangement during collective migration. The local modulus *G* is equal to (1) *G* ≡ *G*_*s*_~*α*_1_*ρ*_*A*_^*β*_1_^ (where *G*_*s*_ is the elastic shear modulus) for the case of shear stress/strain and (2) *G* ≡ *E*~*α*_2_*ρ*_*A*_^*β*_2_^ (where *E* is Young’s modulus) for the case of normal stress/strain. The Young’s modulus obtained for epithelium is *E* = 1 *kPa* (39).

The number density of cell adhesion contacts can be expressed as *ρ*_*A*_ = ∑_*l*_ *δ*(ℜ − *r*_*l*_). The corresponding free energy density of adhesion contacts can be expressed as *F*_*A*_ = ∑_*l*_ *μ*_*l*_*δ*(ℜ − *r*_*l*_) (where *μ*_*l*_ is the chemical potential of the l-th adhesion contact). The density of adhesion contacts changes as the product of (1) biochemical processes such as cell signalling and gene expression and (2) accumulation of strain energy caused by moving through a dense environment. Consequently, the cumulative change of the density of adhesion contacts *ρ*_*Aj*_(ℜ, *τ*) (where *j* can be *j* ≡ *m* for migrating cell cluster and *j* ≡ *BL* for surrounding boundary layer) can be expressed thermodynamically by formulating a Langevin-type equation:

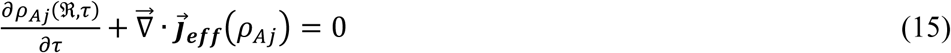

where 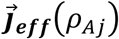 is the effective flux equal to

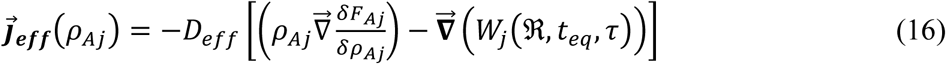

where *W*_*j*_(ℜ, *t*_*eq*_, *τ*) is the local strain energy density within a migrating cluster or surrounding boundary layer.

We introduced the following assumption in order to simplify further modeling consideration:

1. Compressive residual stress within migrating cluster is significantly larger than shear stress. The resistance of migrating cluster to shear stress can be regulated by the ratio 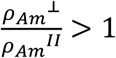 (where *ρ*_*Am*_^⊥^ is the density of adhesion contacts perpendicular to the direction of migration, *ρ*_*Am*_^*II*^ is the density of adhesion contacts parallel to the direction of migration).
2. Shear stress within perturbed boundary layer is significantly larger than normal stress such that 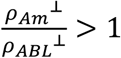 (where *ρ*_*ABL*_^⊥^ is the density of adhesion contacts within the boundary layer perpendicular to the direction of migration).

### 2.2 Averaged shear stress generated within a perturbed boundary layer

Step displacements of the migrating cell cluster induce the generation of local compressive and shear strains within migrating cluster and within the perturbed boundary layer. These strains lead to a generation of local stresses and their relaxations during successive relaxation cycles (4). Schematic representation of the successive relaxation cycles is shown in Figure 2.

**Figure 2.**
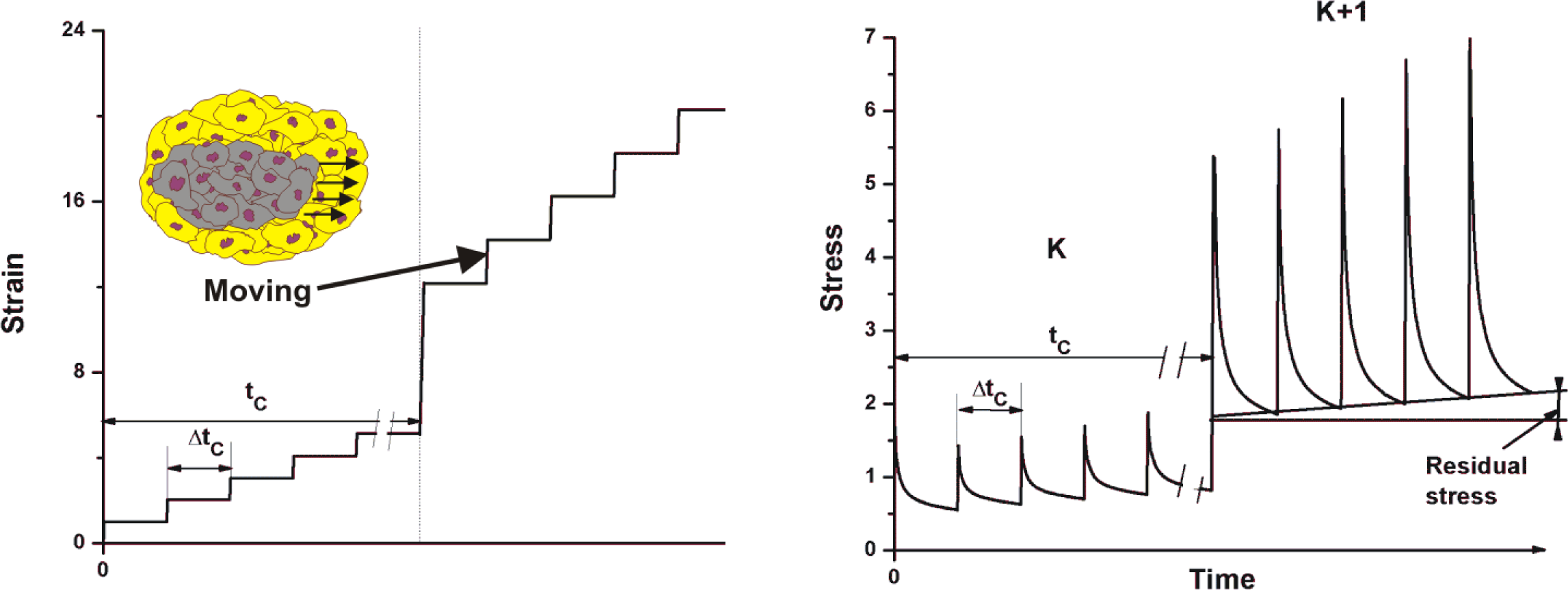
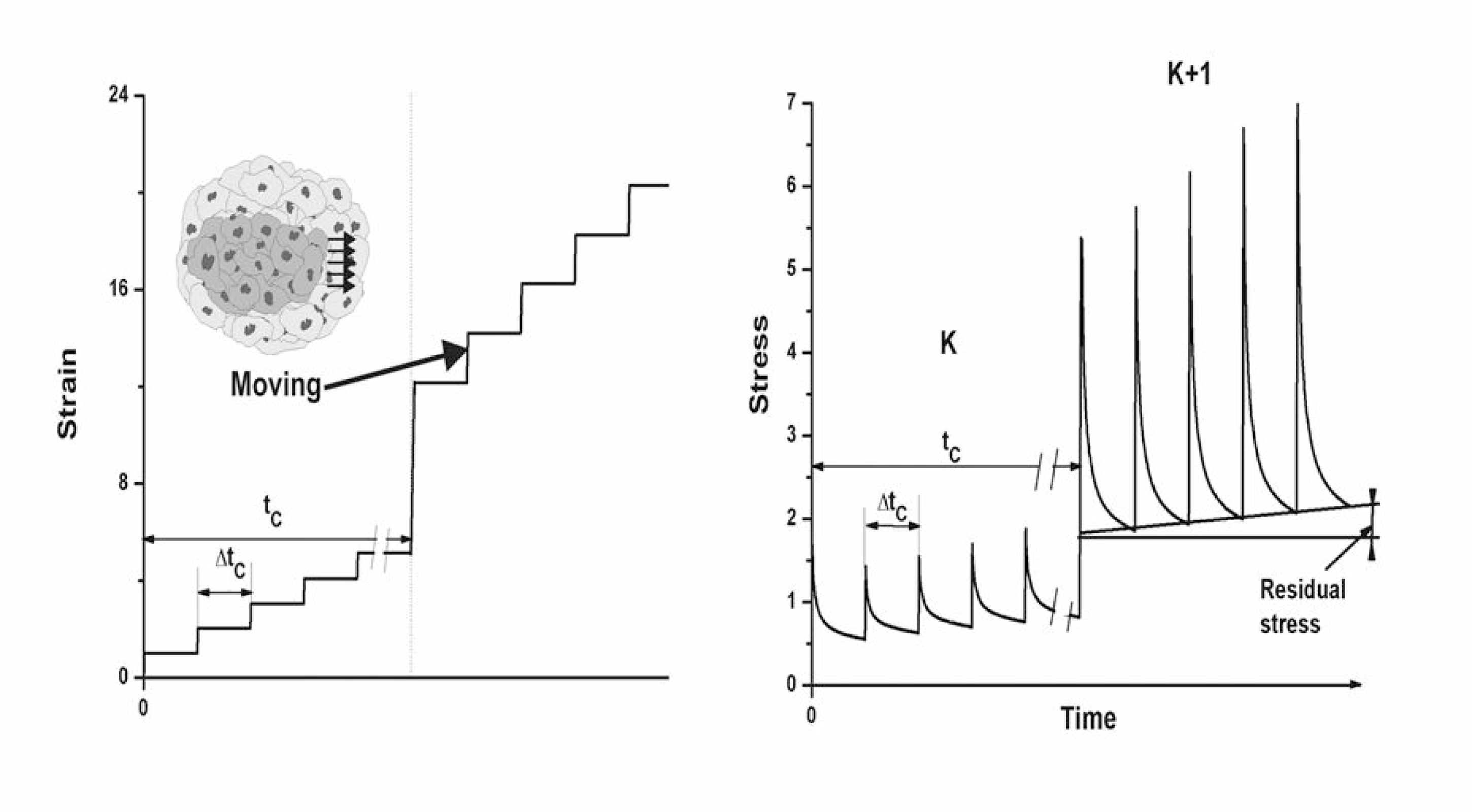
Schematic representation of successive relaxation cycles caused by the movement of cell cluster through dense surrounding made by resting cells; Interactions of two viscoelastic systems such as migrating cluster and surrounding perturbed boundary layer induces multi-time relaxation cycles.

We would like to provide the rough estimation of the magnitude of average viscous part of shear stress within the perturbed boundary layer *σ*_*BL Svis*_ depending on the layer thickness and effective viscosity. The average viscous component of shear stress within the boundary layer around the migrating cell cluster can be expressed as:

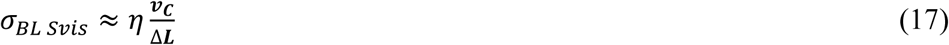

where ∆*L* is the thickness of the perturbed boundary layer, *v*_*C*_ is the speed of migrating cell cluster and *η* is the viscosity. Marmottant et al. (33) determined the effective viscosity of multicellular epithelial aggregate equal to *η* = 4.4*x*10^5^*Pas*. Clark and Vignjajevic (28) reported that the speed of migrating cell clusters during embryogenesis is about 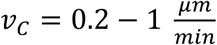. We supposed that the bilayer thickness could be the order of magnitude higher than the size of single cell, i.e. ∆*L* ≈ 100 *μm*. Corresponding values for the shear stress component is in the range of 15-73 *Pa*. This range of shear stress corresponds to the experimental data reported by Tambe et al. (8). The value of shear stress can be reduced by (1) increasing the thickness of the boundary layer ∆*L* or (2) decreasing the effective viscosity *η*. The value of the effective viscosity depends on the number density and type of adhesion contacts within a boundary layer, i.e. *η*~*ρ*_*A BL*_.

### 2.3 Stress accumulation during the collision of migrating cell clusters

Collisions of velocity fronts are a consequence of uncorrelated motility primarily caused by time delay in cell response to various mechanical and biochemical stimuli. The delay effects are induced by gene expression (6). Often, one gene regulator controls another, and so on, in a gene regulatory network. Post-translational modification of membrane proteins, such as phosphorylation and glycosylation, may only require a few minutes, whereas synthesis of proteins and their transport can take tens of minutes and influence short-time relaxation cycles (6).

The strain energy density of migrating cell cluster at the biointerface is equal to ∆*W*_*m*0_(ℜ_*i*_) and formulated by eq. 1. After a collision of two migrating cell clusters, the strain energy density ∆*W*_*m*_(ℜ_*i*_) locally increases i.e. ∆*W*_*m*_(ℜ_*i*_) → ∆*W*_*m*0_(ℜ_*i*_) + *W*′ (where *W*′ is the perturbed energy part). If the local energy density of migrating cell cluster after collision reaches the threshold value *W*_*c*_, cell migration can be suppressed. This threshold value has not been experimentally estimated yet. This is the necessary condition for a formation of the stagnant zone as have discussed qualitatively by Notbohm et al. (9). Nnetu et al. (10) experimentally considered collision of two velocity fronts which suppresses cell migration.

For a deeper understanding of configuration changes of migrating cells, it is necessary to: (1) estimate stress relaxation during 3D collective cell migration, (2) correlate the stress relaxation ability with type and distribution of adhesion contacts within both subsystems (migrating clusters and surrounding perturbed boundary layers), and (3) estimate the threshold value of the strain energy density *W*_*c*_. It is difficult to measure these parameters during 3D collective cell migration under *in vivo* conditions. Consequently, we tried to elaborate cell responses under various stress conditions based on literature.

## 3. Results and discussion

Movement of migrating cell cluster through surrounding tissue can be considered as interactions between two viscoelastic subsystems, i.e. the cluster and the perturbed boundary layer. These interactions induce many short-time relaxation cycles during long-time relaxation cycle which lead to: (1) oscillatory change of normal and shear stresses within both subsystems and (2) accumulation of normal and shear residual stresses within them (4,9). Cell response can be estimated in the context of characteristic times. The stress relaxation time of 3-14 *min* (33) is the same order of magnitude as the duration of short-time relaxation cycle is ∆*t* ≈ 10 − 50 *min* estimated by Pajic-Lijakovic and Milivojevic (4) while the duration of long-time relaxation cycle corresponds to a few hours (7,21). Tambe et al. (8) pointed out that 2D collective migration of epithelial cells results in the generation of normal and shear stresses up to 100-150 *Pa*. It can be expectable that stress accumulation is more intensive in 3D. We estimated the average value of the viscous part of shear stress generated within a perturbed boundary layer in the range of 15-73 *Pa* calculated for the layer thickness equal to ∆*L* ≈ 100 *μm* and effective viscosity *η* = 4.4*x*10^5^*Pas* determined by Marmottant et al. (33).

Cells are more sensitive to the action of shear stress compared with the compressive stress. Shear stress higher than a few *Pa* significantly influences the state of epithelial cells (15). However, cells well tolerate compressive stress up to a few *kPa* (17,18). Hampel et al. (40) considered the response of corneal epithelial cell monolayers under (1) constant shear stress equal to 1.4 *Pa* and (2) a high shear stress oscillatory profile with a peak of 4 *Pa* during 2 h. Gene expression of E cadherin and tight junction protein 1B under oscillatory shear stress is increased compared with constant shear stress. Molladavoodi et al. (41) applied the shear stress of 0.4 to 0.8 *Pa* to estimate the response of corneal epithelial cell monolayer and pointed to the time-dependence of cytoskeletal rearrangement. Flitney et al. (15) reported that some cells, most notably by endothelial cells well tolerate shear stress up to 1.5 *Pa*. On the other hand, higher shear stress especially when sustained for long periods, can be detrimental and may result in severe damage to vimentin and keratin intermediate filament networks. Pitenis et al. (16) pointed out that shear stress of ~60 *Pa* is sufficient to induce inflammation in epithelial cells during 5.5 h. Partial disintegration of TJs can contribute to inflammatory effects (42). Considered values of shear stress are much lower than the stress generated at single AJ between two contractile cells. Liu et al. (14) reported that the stress at single AJ between two active cells is approximately constant and equal to 1 *kPa*.

On the contrary, much higher compressive stress has been applied in the literature in order to estimate cell response. Srivastava et al. (43) considered cell migration under mechanical compression using Dictyostelium cells as a model. They reported that mechanical compression of the order of 500 *Pa* directly triggers a transition in the mode of migration from primarily pseudopodial to bleb driven in <30 s. Cheng et al. (17) considered the influence of compressive mechanical stress produced within 0.5% agarose hydrogel matrix during murine breast carcinoma cell line EMT6 tumor spheroid growth on cell migration and proliferation. They pointed out that compressive stress of ~780 *Pa* significantly influences cell migration. Tse et al. (18) reported that highly aggressive 4T1 and MDA-MB-231 cells, as well as 67NR cells stopped proliferating and started to create a leader cell formation, which allowed them to move. Partial epithelial–mesenchymal transition (EMT) is provoked by applied compressive stress of ~600 *Pa*. TJs disappear during EMT (44). They pointed out that TJs are vital to the prevention of cancer cell metastasis. Many questions arise. How can cells adapt to the action of higher shear stress and how can cells minimize the shear stress generation?

Single cells can adapt to the action of shear stress by phosphorylation of keratin intermediate filament network (KIF). The process of adaptation depends on the magnitude of stress and time of cell exposure (15). Phosphorylation requires a few minutes, whereas the synthesis of proteins and their transport can take tens of minutes (6). Consequently, the concentration of KIF can be increased in order to protect cells against shear stress during several tens of minutes (45). The phosphorylation is intensive in the region of KIF near nucleus under a modest shear stress of 1.5 *Pa* during 1 h (15). However, we should consider here cell response under shear stress up to 100-150 Pa.

In the next step of this consideration, we discussed the possible ways of cells within migrating clusters and perturbed boundary layers to minimize the undesirable generation of shear stress. Migrating cell clusters could reinforce their structures by forming stronger adhesion contacts such as tight junctions (TJs) perpendicular to the direction of movement and protect themselves against shear stress. On the other hand, larger number density of adherens junctions (AJs) parallel to the direction of migration enables the elastic stretching of the cluster in the direction of migration which reduces friction effects during its movement (46). AJs promote a relatively weak cell–cell adhesion compared with desmosomes or TJs (47). The phenomenon can be discussed in the context of the ratio of the number densities of TJs relative to AJs:

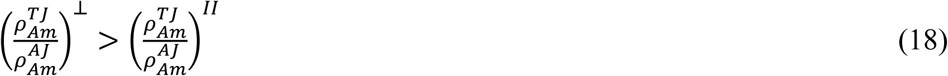

where 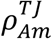 is the number density of TJs and 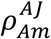 is the number density of AJs perpendicular “⊥” and parallel “*II*” to the direction of migration. The lower ratio 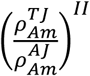 formed parallel to the direction of migration enables the elastic stretching of the cluster in the direction of migration which reduces friction effects during its movement. Shi et al. (47) pointed out to specific role of TJs in regulation of collective cell migration during wound healing. These distributions of adhesion contacts influence cumulative effects of cell signalling through adhesion-mediated signalling pathways involved in actin organization, cell polarity, as well as transcriptional regulation (20,48). The distribution of adhesion contacts per single migrating cluster and its influence to its stiffness inhomogeneity haven’t been considered experimentally yet.

Shear stress generation is more intensive within boundary layers around the migrating cell clusters in comparison with migrating cluster itself. The magnitude of shear stress could be minimized by: (1) decreasing the speed of migrating cell clusters, (2) increasing the thickness of perturbed layers, and (3) reducing the slip effects as well as the effective viscosity within the perturbed boundary layers by decreasing the density of adhesion contacts *ρ*_*A BL*_ and changing of their states under stress. Decrease in the density of adhesion contacts *ρ*_*A BL*_ under stress and perturbation of cell signalling could lead to local circular single-cell movement within perturbed layers in order to (1) decrease sliding and on that base to decrease the effective viscosity and (2) ensure continuity of the multicellular system. This circular movement of cells within perturbed boundary layers has been discussed by Shellard and Mayor (25). Schematic representation of migrating cell cluster and perturbed boundary layer are shown in Figure 3.

**Figure 3.**
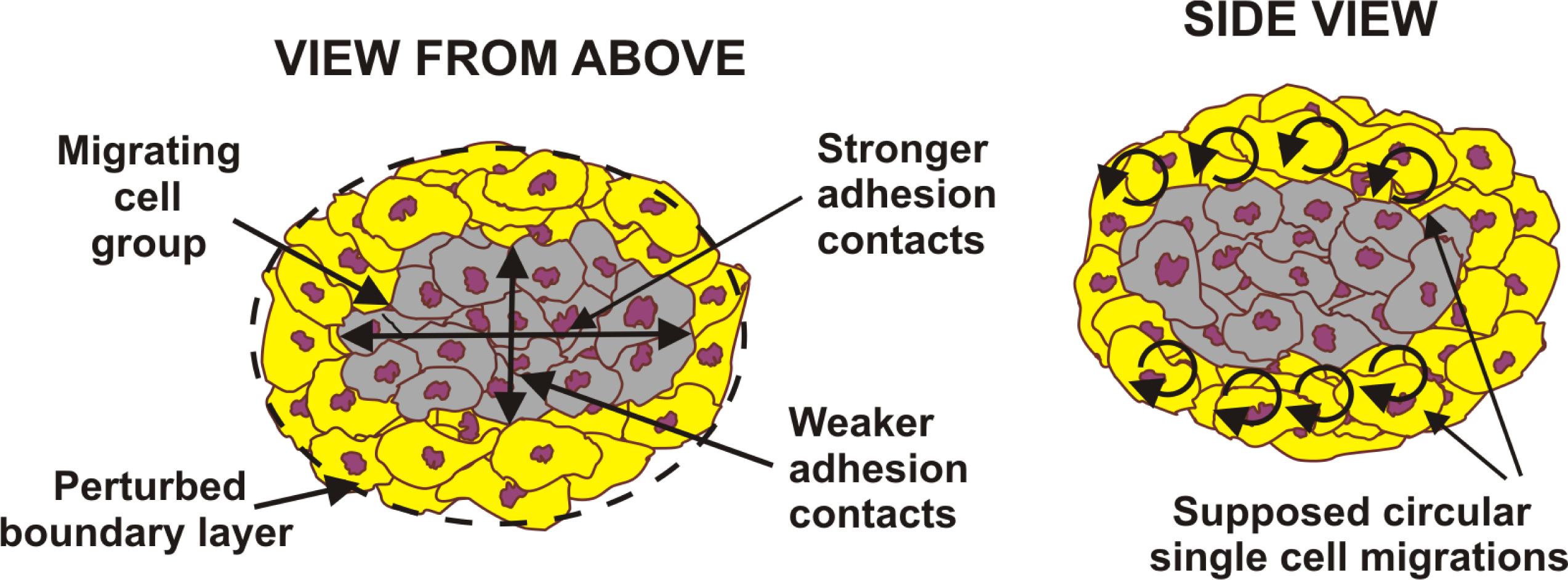
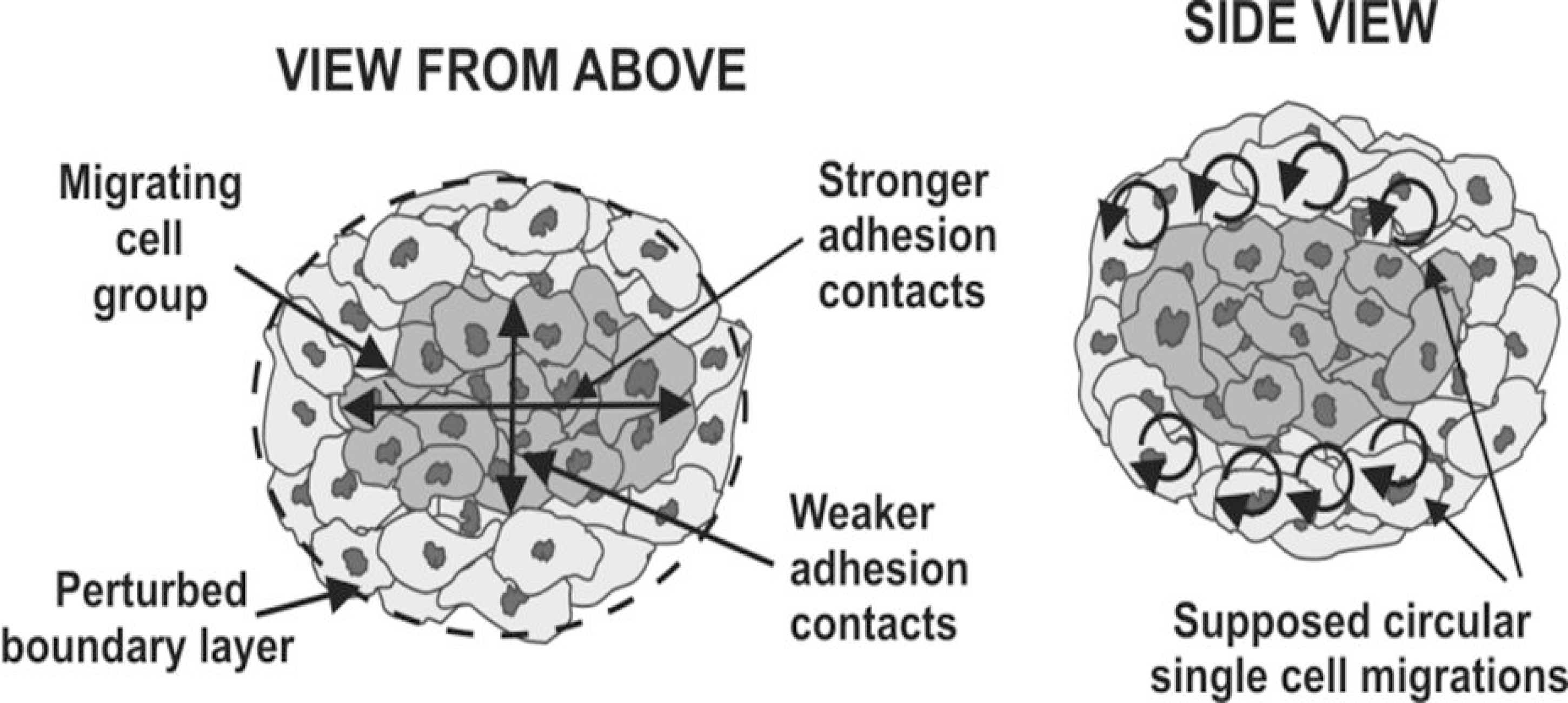
Types of adhesion contacts, their state under stress and the distribution of the cohesiveness density can reinforce migrating cluster against shear stress and influence the state of the perturbed boundary layer.

Additional experiments are needed in order to correlate stiffness distribution with the distribution of TJs per single migrating clusters and perturbed boundary layers.

## 4. Conclusions

Collective cell migration induces a local generation of stress (normal and shear) significant even in 2D (up to *100-150 Pa*). Cells well tolerate compressive stress up to a few *kPa*. However, shear stress of a few *Pa* can induce severe damage to vimentin and keratin intermediate filament networks during 1 h, while shear stress of ~60 *Pa* influences gene expression which can cause the inflammation in epithelial cells during 5.5 h. Cell strategy to minimize undesirable shear stress generation was discussed based on rheological consideration.

Migrating cell clusters can reinforce their structures by forming stronger adhesion contacts such as TJs perpendicular to the direction of movement and protect themselves against shear stress in comparison with type and density of adhesion contacts parallel to the direction of movement. These conditions lead to the establishing of stiffness inhomogeneity per single migrating clusters. The magnitude of shear stress generated within the perturbed boundary layers around migrating clusters can be minimized by: (1) decreasing the speed of migrating cell clusters, (2) increasing the thickness of perturbed layers, and (3) reducing the slip effects as well as the effective viscosity within the perturbed boundary layers by decreasing the density of AJs and TJs as well as changing of their states under stress. The role of TJs in collective cell migration should be clarified from subcellular to supracellular levels.

Additional experiments are necessary to determine stress distribution during 3D collective cell migration and its change and correlate: (1) the stiffness inhomogeneity per single migrating clusters with the type and distribution of AJs and TJs, (2) slip effects within perturbed boundary layers with the type and distribution of AJs and TJs within them, and (3) lifetime of migrating clusters and its distribution with the residual stress distribution within a multicellular system.

## Author contributions

All authors contributed equally to the paper.

## Acknowledgment

This research was funded by grant III46001 from the Ministry of Education, Science and Technological Development, Republic of Serbia.

